# A push-pull system of repressors matches levels of glucose transporters to extracellular glucose in budding yeast

**DOI:** 10.1101/2021.04.20.440667

**Authors:** Luis Fernando Montaño-Gutierrez, Marc Sturrock, Iseabail Farquhar, Kevin Correia, Vahid Shahrezaei, Peter S. Swain

## Abstract

A common cellular task is to match gene expression dynamically to a range of concentrations of a regulatory molecule. Studying glucose transport in budding yeast, we determine mechanistically how such matching occurs for seven hexose transporters. By combining time-lapse microscopy with mathematical modelling, we find that levels of transporters are history-dependent and are regulated by a push-pull system comprising two types of repressors. Repression by these two types varies with glucose in opposite ways, and not only matches the expression of transporters by their affinity to a range of glucose concentrations, but also the expression of some to how glucose is changing. We argue that matching is favoured by a rate-affinity trade-off and that the regulatory system allows yeast to import glucose rapidly enough to starve competitors. Matching expression to a pattern of input is fundamental, and we believe that push-pull repression is widespread.

## Introduction

Gene expression is regulated so that genes are expressed opportunely [1]. The presence of some regulatory molecules and the absence of others allows transcription to initiate. A more complex but still common requirement is for a gene to be expressed only within a range of concentrations of a regulator. A well known example occurs during the development of the fly embryo where some zygotic genes are expressed only if their regulatory transcription factors have concentrations that are neither too low nor too high [2]. Less recognised is that similar regulation occurs in unicellular organisms, whose genomes often encode multiple transporters for the same nutrient [3].

The model eukaryote budding yeast, for example, has 18 transporters for glucose [4]. Seven of these are necessary for growth [5], and where investigated each transporter is expressed within a specific range of glucose concentrations [6, 7, 8, 9, 10].

Using yeast’s glucose transport as an exemplar, we set out to understand mechanistically how gene expression is matched to a range of a regulatory molecule. At the same time, we wished to better understand why yeast has evolved so many transporters and why restricting their expression is advantageous. Our approach was to generate time-series data. We fluorescently tagged the transporter genes and quantified the levels of the transporters in bulk and in single cells in dynamically changing extracellular glucose, both for the wild-type and for strains with a perturbed regulatory network. To integrate these diverse data, we used mathematical modelling and statistical inference.

Yeast’s glucose transporters are hexose transporters and are encoded by genes HXT1 through HXT17 [4] and by GAL2, a transporter for galactose that also has a high affinity for glucose [11]. At least one of HXT1 through to HXT7 is needed for growth on glucose [5]. Eight of the other HXT genes are located near the end of chromosomes and their high number may be because of genomic drift, generated inadvertently as the cell maintains its telomeres [12]. The network regulating the HXT genes has mostly been identified [4, 13] and is embedded within the cell’s larger signalling network that coordinates growth with the availability of nutrients [14].

We begin by showing that Hxts should exhibit a rate-affinity tradeoff [15] because they use facilitated diffusion [16]. This tradeoff, together with each Hxt having a unique affinity for glucose [11, 17], implies that there is a range of glucose concentrations for which an Hxt imports glucose at a greater rate than any other. Although behaving broadly consistently with our observations, cells express not only the optimal Hxt but also several others. We interpret this behaviour as cells preparing for a change in glucose’s availability. Using microfluidics and time-lapse microscopy, we demonstrate that for certain inputs the levels of Hxts can be history-dependent: some increase only in rising glucose and others only in falling glucose. Extending this data to mutants of the regulatory network and complementing it with broader studies in batch culture, we develop a quantitative, predictive understanding of the regulation, which not only matches expression to a range of glucose concentrations but also to how glucose is changing. The dynamics of almost all seven Hxts is generated by four repressors that interchange at the HXT promoters in a push-pull manner as glucose alters, in a way that is analogous to behaviour in *Drosophila* embryos. Finally, we argue that the regulation and number of HXTs make budding yeast excel at rapidly importing glucose compared to other fungi.

## Results

### A rate-affinity tradeoff predicts a range of expression for each HXT

Of the seven transporters necessary for growth on glucose [5], each has a different affinity for glucose [17, 11]: HXT1 and HXT3 have a low affinity; HXT2 and HXT4 have a medium affinity; and HXT6 and HXT7 have a high affinity (Fig. 1A). HXT5 has medium affinity too, but its regulation is thought to be distinct [19, 20].

**Figure 1.**
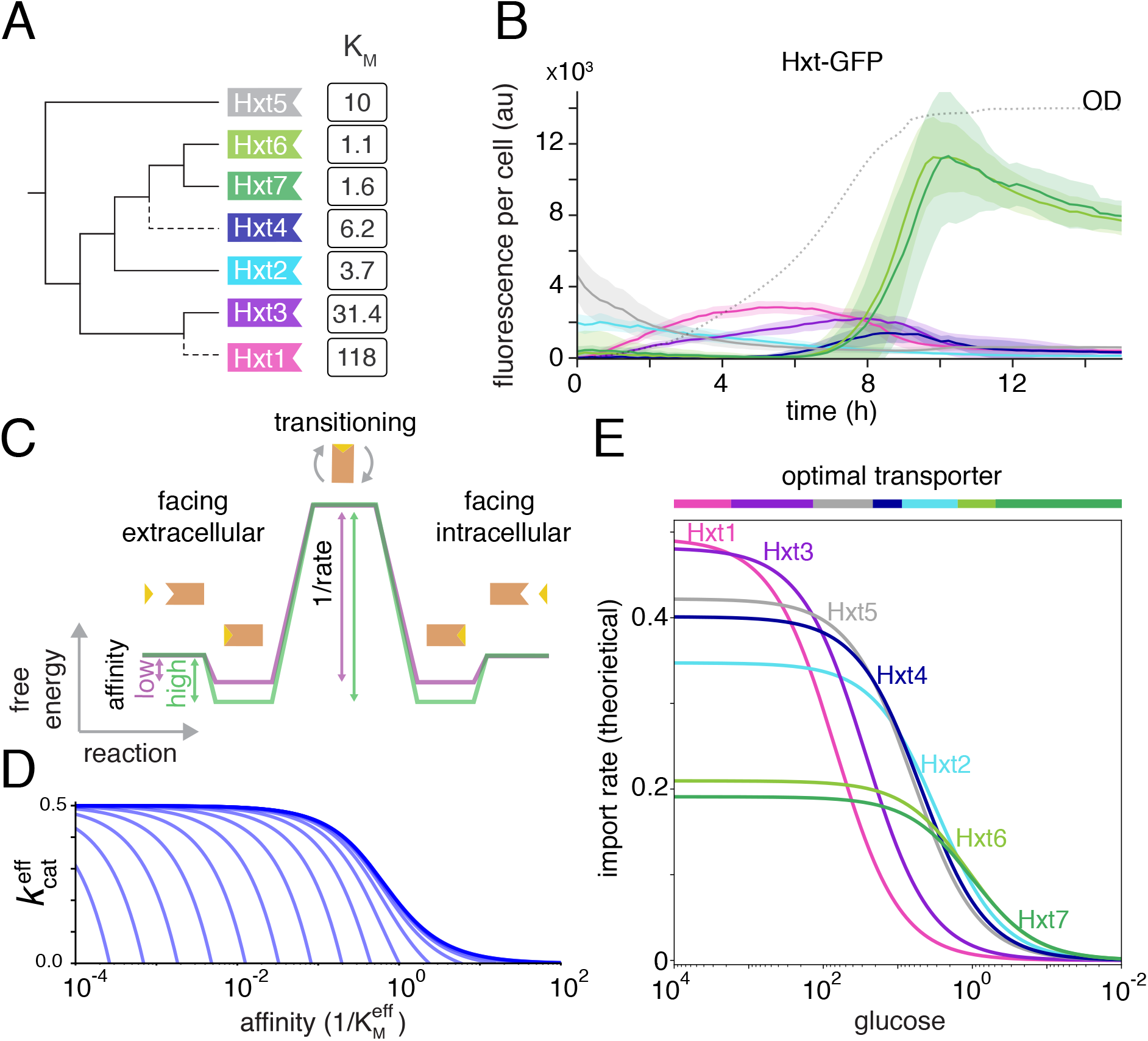
A rate-affinity tradeoff explains why budding yeast has multiple different hexose transporters. **A** Budding yeast has seven hexose transporters of which at least one is necessary for growth on glucose. Each transporter has a different affinity given by the inverse of its *K_M_* (here in mM). We show the transporters’ phylogenetic relationships (SI), with dashed lines indicating the whole genome duplication [18]. **B** The different HXTs are expressed in an order approximately determined by their *K_M_* as glucose falls (colours as in **A**). We follow transporters tagged with GFP in batch culture with initially 2% (110 mM) glucose. The concentration of glucose decreases as the culture’s optical density (OD) increases (dotted line) and is near zero when the OD plateaus. **C** The rate-affinity tradeoff may be understood from a reaction coordinate diagram. High affinity transporters (green) necessarily have a lower rate than low affinity transporters (purple). Glucose is depicted as a triangle and the transporter as an orange polygon. **D** Mathematically modelling facilitated diffusion, we can calculate the rate-affinity tradeoff for an effective rate and affinity that depend on intracellular glucose. Each line corresponds to a different concentration of intracellular glucose with this concentration decreasing from the origin and being zero at the bold line. **E** We predict that each transporter has a range of glucose concentrations where it generates the highest flux of imported glucose (upper bar – assuming negligible intracellular glucose for simplicity). The rate-affinity tradeoff implies that for a particular glucose concentration one type of Hxt has the highest flux.

During growth in batch culture, the transporters are mostly expressed sequentially in time following their affinity (Fig. 1B). Tagging the HXT genes with Green Fluorescent Protein (GFP), we observed that, for cells in 2% glucose, the low affinity transporters – Hxt1 and then Hxt3 – are the first to peak when the concentration of glucose is highest, followed by the medium affinity transporter Hxt4, and then the high affinity transporters Hxt6 and Hxt7, which peak as glucose is exhausted. The medium affinity transporters Hxt2 and Hxt5 do not fit this pattern. The level of Hxt5, in particular, decreases monotonically in glucose [21].

This sequential expression is consistent with the transporters having a rate-affinity tradeoff (SI). The Hxts transport using facilitated diffusion [16]. Embedded in the plasma membrane, their structures fluctuate – facing inwards towards the cytosol, then outwards towards the extracellular space, and then back again – and so they passively transport glucose from high to low concentrations. The potential for a trade-off may be understood intuitively using a reaction coordinate diagram (Fig. 1C) [22]. A transporter’s affinity is determined by the difference in free energy between glucose in solution and glucose bound to the transporter – the larger the free energy difference, the higher is the affinity. A transporter’s rate of import is mostly determined by the difference in free energy between the complex with glucose and the transition state as the transporter changes to face the intracellular space – the larger the free energy difference, the lower is the import rate because the activation barrier is greater. We assume that the time taken to cross this barrier is substantially longer than the time taken for glucose to unbind from the transporter and enter the cytosol. An increase in affinity therefore necessarily decreases the rate.

Mathematically modelling facilitated diffusion supports this intuition, although with added complexity because the transporters potentially transport glucose both into and out of the cell [23, 24]. For symmetric transporters, the flux of imported glucose is a Michaelis-Menten function of the difference between the extra- and intracellular glucose concentrations and has an effective *k*_cat_ and *K_M_* that depend on intracellular glucose (SI). Higher intracellular glucose undermines import by both decreasing the effective *k*_cat_ and increasing *K_M_* (Fig. 1D).

The tradeoff implies that for a given concentration of extracellular glucose there is a particular transporter, defined by its effective *K_M_*, that maximises the flux of imported glucose. A transporter with a lower effective *K_M_* generates less flux because of the corresponding decrease in its effective *k*_cat_; a transporter with a higher *K_M_* generates less flux because of its lower affinity (SI). Using the measured *K_M_* values (Fig. 1A) [17, 11], we predicted the order in which the transporters should be expressed if the cell is to maximise its import of glucose (Fig. 1E). This prediction relies on strong assumptions – that there is a fixed amount of cellular resources dedicated to hexose transporters, negligible free intracellular glucose, and that steady-state behaviour is reached quickly – yet the predicted order is broadly consistent with the peak of each transporter’s expression as glucose decreases (Fig. 1B).

A rate-affinity tradeoff also implies that the cell should express the HXTs one at a time to maximise the import of glucose. It is always best to put the available intracellular resource into synthesising the transporter optimal for a given concentration of extracellular glucose than to share that resource between optimal and suboptimal transporters.

One-at-a-time expression is not, however, what we observed (Fig. 1B). Including rates of synthesis and degradation of the transporters in the mathematical model shows that co-expressing two Hxts – the optimal transporter and the transporter closest to being optimal – may increase the overall steady-state flux (SI). If transporters are synthesised from a fixed, common pool of precursors, such as amino acids, and replenish that pool when degraded, then this increase in overall flux arises because more of the precursors are put into use when two transporters are expressed. With two transporters, the total rate of synthesis is higher than with only one transporter. Although expressing the second transporter reduces the amount of optimal transporters and so lowers the flux they generate, this loss can be more than compensated by the flux through the suboptimal transporters – if both their numbers are sufficiently high and the flux through each is sufficiently close to the optimal transporter’s flux.

Yet the cells’ response is richer even than this prediction because more than two Hxts are co-expressed (Fig. 1B), although in batch culture which is not at steady state. The level of intracellular resources also appears not to be fixed because there is substantially higher expression of transporters when glucose is scarce compared to when it is plentiful (Fig. 1B).

To investigate simpler and steady-state-like behaviours, we used microfluidics and time-lapse microscopy [25, 26, 27], where cells grow in a constant flow of medium with a controlled concentration of glucose.

### Levels of the Hxts are history-dependent

After 10 hours of growth in constant glucose, we find that multiple transporters are still coexpressed for all concentrations of glucose tested (Fig. 2A). The affinities of the mostly highly expressed transporters match the glucose concentration – Hxt6 and Hxt7 in low 0.01% glucose; Hxt2 in medium 0.1% glucose; and Hxt1 and Hxt3 in high 1% glucose. As expected from our modelling, the co-expressed transporters have similar affinities to the optimal transporter, but the data are inconsistent with cells regulating expression to maximise their import for the current concentration of glucose.

**Figure 2.**
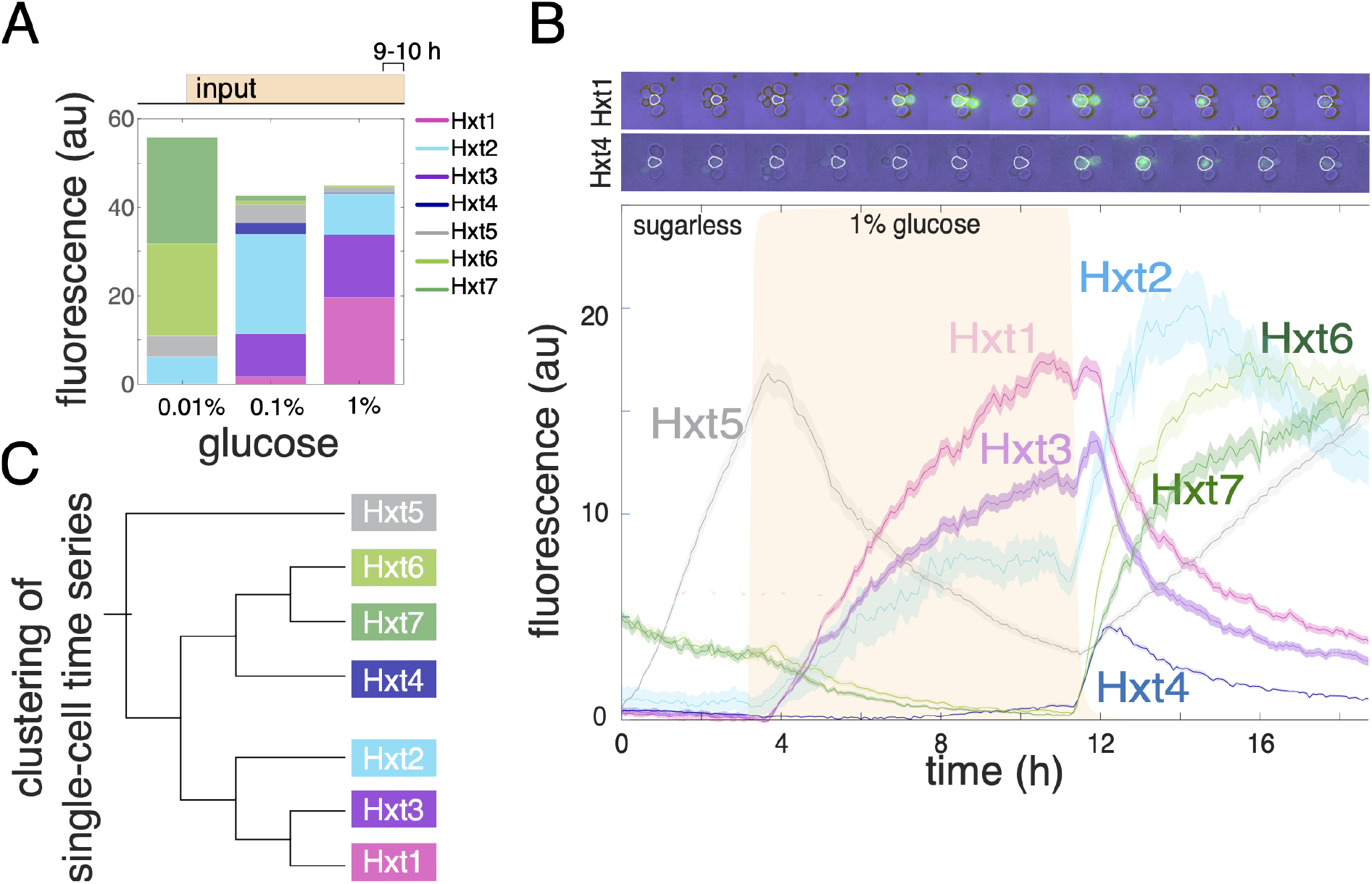
In changing glucose, levels of Hxts are history-dependent. **A** After 10 hours in constant concentrations of glucose, not one but multiple Hxts are present. Except for Hxt2, only transporters with similar values of *K_M_* are co-expressed. We show fluorescence for GFP-tagged Hxts averaged over the last hour of each experiment. **B** In changing glucose concentrations, multiple HXTs express simultaneously and their levels are history-dependent. Some even respond only to falling concentrations of glucose. Top: time-lapse microscopy images of cells in a single trap of the ALCATRAS microfluidic device [25] showing GFP-tagged Hxts. The central cell is outlined in white. Bottom: The mean levels of Hxts shown as a function of time for cells growing in SC medium where glucose rises from zero to a concentration of 1% at approximately 3 hours and falls to zero again at approximately 11 hours. Errors are standard deviations over at least two experiments. **C** Clustering the single-cell time series from **B** shows the similarities in the mean behaviours of the different Hxts is maintained at the single-cell level and largely reproduces the phylogenetic relationships of Fig. 1A.

Rather our results suggest that cells regulate the Hxts to satisfy two goals: both to have high import of the glucose currently available and also to prepare for possibly impending changes in the concentration of glucose [28, 29].

To test this hypothesis, we submitted cells to a rising, constant, and then falling glucose concentration in a dynamic but controlled environment, switching from a sugarless medium into eight hours of a constant glucose and back to the sugarless medium.

We observed a remarkable diversity of behaviour (Fig. 2B). As expected if cells prepare for likely future levels of glucose, there is a clear dependence on history, with the response differing if glucose is rising or falling. Levels of the medium affinity transporter Hxt4 and the low affinity transporters Hxt6 and Hxt7 spike but only in decreasing glucose; levels of the medium affinity transporter Hxt2 spike when glucose both starts to increase and to decrease; levels of Hxt5 appear to increase exclusively in the absence of glucose, consistent with earlier work [30]; and only the low affinity transporters Hxt1 and Hxt3 behave as conventionally expected – their levels increase if and only if the glucose concentration is sufficiently high.

These behaviours hold too at the single-cell level and there are no detectable sub-populations. Using mutual information to cluster the single-cell time series [31] (SI), the clusters match those anticipated from the mean behaviour (Figs. 2C & 2B) and, excepting Hxt2, that expected from phylogeny (Figs. 2C & 1A).

Given these history-dependent responses, we asked if they are consistent with the known regulation of the HXT genes.

### The Hxts are a focal point of regulation

Appropriately, considering glucose’s central metabolic role, regulation of the HXTs is complex. There are three main contributors [14]: the Snf3-Rgt2 network that directly senses extracellular glucose; SNF1 kinase – a complex of three proteins and known as AMP kinase in higher eukaryotes, which promotes growth on other sources of carbon as glucose falls; and protein kinase A, which regulates the biogenesis of ribosomes and promotes growth if glucose rises.

The Snf3-Rgt2 network comprises a low affinity sensor for glucose – Rgt2, a high affinity sensor – Snf3, and a transcriptional regulator, Rgt1, together with its two co-repressors Mth1 and Std1 [32] (Fig. 3A). If Rgt1 forms a complex with at least one of Mth1 and Std1 [33], it recruits the general repression machinery to the HXT promoters and inhibits expression [6, 34]. In the presence of glucose, however, Snf3 or Rgt2 or both cause the co-repressors Mth1 and Std1 to inactivate [35], enabling the HXTs to express. Mth1 is inactivated by degradation [36]. Although both Mth1 and Std1 are actively degraded in glucose, only levels of Mth1 decrease because of compensatory expression from the STD1 gene [37, 38]. Std1 is inactivated by forming condensates outside of the nucleus if the glucose concentration is sufficiently high [39] (Fig. 3B).

**Figure 3.**
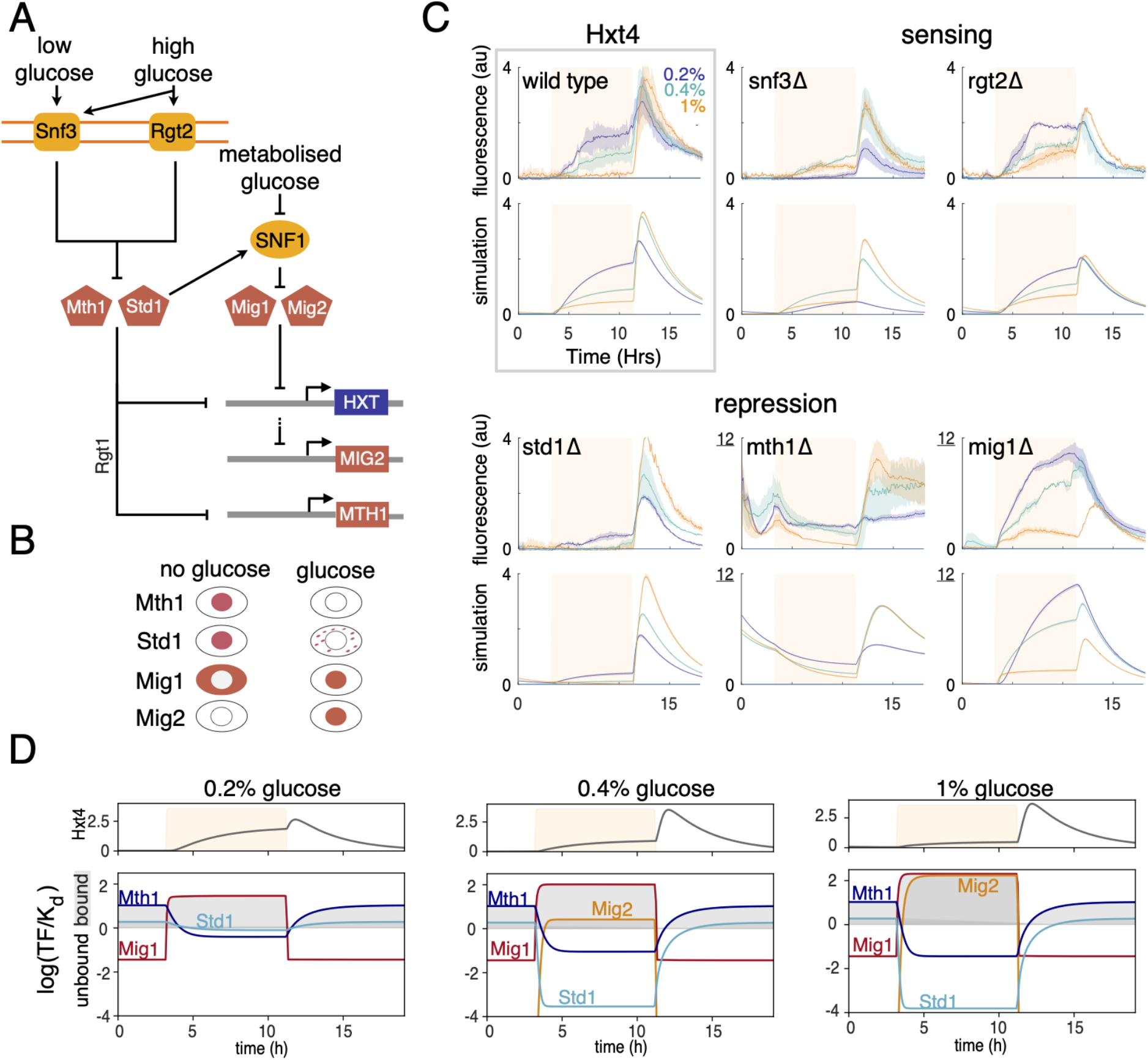
Levels of Hxt4 are controlled by repressors Mth1 and Std1 interchanging with repressors Mig1 and Mig2 at the HXT4 promoter. **A** HXT genes are regulated by four repressors. Mth1 and Std1 repress when bound with the transcription factor Rgt1 at the HXT promoters and are inactivated by the sensors Snf3 and Rgt2. Mig1 and Mig2 are inactivated by SNF1, which is activated by low levels of metabolised glucose and by interacting with Std1. The MIG2 gene is repressed by Mth1 and Std1; MTH1 is repressed by Mig1 and Mig2; and STD1 by Rtg1 (not shown because levels of Std1 are constant as glucose changes). **B** This regulatory network causes Mth1 and Std1 to be in the nucleus (inner circle) and repressing in sufficiently low glucose. In contrast, Mig1 and Mig2 are in the nucleus and repressing in sufficiently high glucose. **C** A systematic microfluidics-based study of Hxt4 enables a quantitative determination of its regulation. We expose cells to dynamic inputs of glucose that rise from zero, remain constant – at a concentration of either 0.2% (purple), 0.4% (cyan), or 1% (orange) – and then fall back to zero. Both the wild-type and strains deleted for a component of the regulatory network are investigated. We fit a mathematical model of the network of **A** – top shows mean fluorescence with the standard deviation across at least two replicate experiments as errors; bottom shows the best-fit model. Notice the different y-axes for the *mth1* Δ and *mig1* Δ strains. **D** Nuclear Mig1 and Mig2 are predicted to respond faster to changing glucose than Mth1 and Std1, which enables a spike of expression from HXT4 in falling glucose. Using our model, we plot the log_2_ of the predicted level of repressor scaled by its dissociation constant of binding to the HXT4 promoter. The transcription factors repress only when these variables are positive. The peak of the spike in Hxt4 levels corresponds with the point when Mth1 starts again to repress.

SNF1 kinase responds indirectly to low intracellular glucose, possibly through the availability of ADP [40], and regulates the HXTs through two repressors Mig1 and Mig2 [37] (Fig. 3A). Active SNF1 promotes HXT expression by both phosphorylating Mig1 [41], which then exits the nucleus (Fig. 3B), and inducing the degradation of Mig2 [42]. The MIG2 gene is regulated by Rgt1 and so expresses in glucose [37]. Mig1 and Mig2 repress the HXTs and also the MTH1 gene for the co-repressor [37].

Furthermore, SNF1 binds the other co-repressor Std1 [43, 44], which also functions as a signalling protein. When bound, Std1 stimulates SNF1’s activity [45], but cannot do so when driven into condensates by glucose [39].

Finally, protein kinase A (PKA) affects HXT expression by phosphorylating Rgt1 when Rgt1 is free of Mth1 and Std1 [46]. This phosphorylation turns Rgt1 into an activator, at least at the HXT1 promoter, but Rgt1 eventually unbinds the DNA when fully phosphorylated by PKA [46, 36].

### Regulation is revealed by the response of Hxt4 to dynamic inputs and genetic perturbations

To understand how this network generates history dependence, we focused on the regulation of Hxt4, which has distinct behaviour in rising and falling glucose (Fig. 2B). Our approach was to apply similar inputs of glucose to mutants missing a gene of the regulatory network and so infer the role played by the deleted gene through comparing the mutant levels of Hxt4 with the wild-type response. For Mig1 and Mig2, we focused on the *mig1* Δ strain because Mig2 typically acts redundantly with Mig1 [47].

Inspecting these data reveals four key points (Fig. 3C):

i. The repressors Mth1 and Std1 are not redundant [48, 38, 49] and differently affect expression of HXT4.
ii. HXT4 is strongly repressed both by Mth1 in the absence of glucose and by Mig1 in the presence of glucose.
iii. As expected, deleting SNF3 – the high affinity sensor – affects the behaviour most in low glucose [50], and deleting RGT2 – the low affinity sensor – affects the behaviour most in high glucose [32].
iv. Counterintuitively, the levels of Hxt4 decrease in the wild-type strain when the constant concentration of glucose midway through the time series increases in value, and this effect is enhanced if the repressor gene STD1 is deleted.

### History dependence is generated by an interchange of repressors

We used mathematical modelling to determine quantitatively whether the data supports the known regulation of HXT4, focusing on the Snf3-Rgt2 network and SNF1 because the regulation by PKA appears mostly to boost already active expression [46]. We assume that Rgt1 is always bound to the HXT4 promoter for simplicity. Although we allow MTH1 to be repressed by Mig1 and Mig2 and MIG2 to be repressed by Std1 and Mth1 [37], we do not model expression of STD1 and MIG1 because their levels are mostly constant [38, 51]. Through the sensors Snf3 and Rgt2, we let glucose inactivate Std1 and degrade Mth1. SNF1 kinase is inactivated by glucose and activated by Std1, and active SNF1 reduces repression by Mig1 and Mig2, causing Mig1 to exit the nucleus and Mig2 to be degraded. Comparing the response of GFP expressed from the HXT4 promoter to GFP-tagged Hxt4 (SI), we allow glucose to protect Hxt4 from degradation [52]. Finally, we assume that the magnitude of all regulation caused by intracellular glucose to be proportional to the level of extracellular glucose – an assumption that subsumes the activities of Hxt4, the other Hxts, and glycolysis. This assumption is retrospectively justified by the accuracy of the fitting we perform.

The model is able to reproduce the behaviour observed (Fig. 3C). We used approximate Bayesian computation to both fit the model to the data and compare models, which showed that there is not strong evidence for a simpler model, lacking some of the regulatory interactions (SI).

The data support HXT4 being controlled by push-pull repression with the four repressors interchanging at the HXT4 promoter when extracellular glucose switches [37] (Fig. 3D). The relevant variables are the nuclear levels of the repressors scaled by their dissociation constants of binding to the HXT4 promoter because only when these variables are greater than one do the repressors actively repress. The repressors work in a push-pull manner. When glucose rises, repression by Mth1 and Std1 decreases – their repression is ‘pulled’ – and is replaced by repression by Mig1 and Mig2 – their repression is ‘pushed’; when glucose falls, the opposite push-pull happens, and repression by Mig1 and Mig2 decreases and is replaced by repression by Mth1 and Std1.

Repression by Mig1 and Mig2 principally determines levels of Hxt4 in rising glucose (Fig. 3D). In a switch into high 1% glucose, repression by Mig1 – our model predicts that Mig1’s repression dominates Mig2’s when both are present (SI) – is both fast and strong enough to prevent any expression even though Mth1 and Std1 are inactivated (Fig. 3D). In a switch into lower glucose (below 1%), there is less Mig1 in the nucleus and levels of Hxt4 do rise. In 0.2% glucose, Std1 activating SNF1 helps reduce Mig1’s repression slightly, but this interaction has a negligible effect at higher glucose concentrations (SI).

The increased repression by Mig2 in higher glucose concentrations (Fig. 3D) is through glucose inactivating Std1 and so de-repressing MIG2 – the inferred parameters are such that Std1 represses the MIG2 promoter substantially more than Mth1 (SI). Although the resulting increased levels of Mig2 little affect HXT4 expression in the wild type because Mig1 dominates and is always present when Mig2 is present, Mig2’s effects are revealed by the mutants. For the *mig1* Δ strain, it is Mig2 that brings down levels of Hxt4 in increasingly higher glucose, and over-expression of MIG2 in the *std1* Δ strain flattens Hxt4’s response.

Levels of Hxt4 spike in falling glucose because of the slow interchange of repression by Mig1,2 to repression by Mth1 and Std1, which creates a window of time where repression is weak and HXT4 expresses. Consistently, the peak in Hxt4 levels for all concentrations of glucose approximately occurs when Mth1 begins again to repress (Fig. 3D).

### Time-series data suggest how the other Hxts are regulated

To investigate how the other Hxts are regulated, we complemented this microfluidics-based work with a broader study performed in batch culture using the same mutants but in strains with one of HXT1-7 fluorescently tagged (Fig. 4). To set the basal level of glucose transport, each strain was pre-cultured in pyruvate where cells perform gluconeogenesis rather than glycolysis before being transferred into glucose.

**Figure 4.**
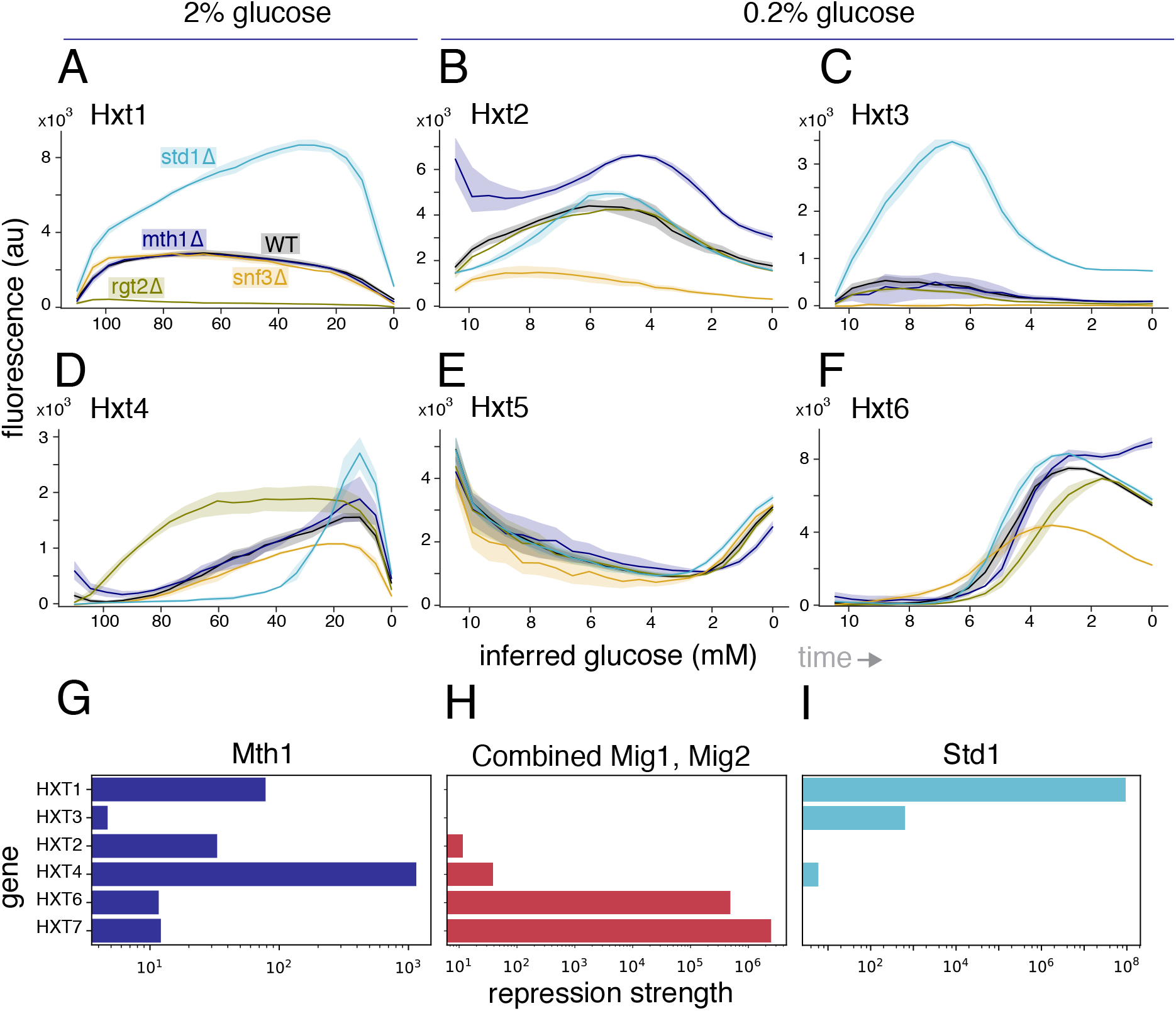
Deviations of HXTs from the wild-type response in strains deleted for regulators suggest particular transcriptional regulation that is supported by mathematical modelling. The wild-type regulation of the Hxts (black) causes most to have their highest levels when their *K_M_* approximately matches the concentration of extracellular glucose (c.f. Fig. 1A). We transformed time-series measurements of GFP-tagged HXTs in batch cultures into functions of extracellular glucose by estimating glucose from the culture’s OD (SI). Time increases from left to right as glucose falls and the OD increases. At the extreme left – corresponding to *t* = 0, the level of Hxt is the level attained in the pyruvate used for pre-growth. We show only representative examples of the full data set (SI). Data are the average of at least two experiments, each with four biological replicates. Errors are 95% confidence intervals. **A** HXT1 is strongly repressed by Std1, and Rgt2 is required to relieve Std1’s repression. **B** HXT2 is strongly repressed by Mth1 but not Std1, and Snf3 is required to relieve Mth1’s repression. **C** HXT3 behaves similarly to HXT1, but expresses in low glucose and requires Snf3 to do so. **D** HXT4 is regulated by Mig1,2 because its levels in high glucose decrease when STD1 is deleted and increase when RGT2 is deleted. **E** HXT5 is only weakly regulated by the network – all mutants behave similarly to the wild-type strain. **F** HXT6 is strongly repressed by Mth1 but not Std1 in low glucose. HXT7 behaves similarly (SI). **G** The predicted strength of repression of Mth1 inferred from the data of Fig. 2B shows that Mth1 represses the medium affinity transporters Hxt2 and Hxt4 more than the high affinity transporters Hxt6 and Hxt7. We define the strength of repression as the maximum of (*R/K_d_*)*^n^* (SI), where R is the time series of the repressor concentration predicted from the regulatory model (Fig. 3B). *K_d_* is the inferred dissociation constant of binding to the promoter and *n* is the inferred cooperativity of binding. **H** The predicted strength of repression of Mig1,2 has the opposite behaviour: stronger repression for the high compared to the medium affinity transporters. **I** Std1 predominately specialise to high extracellular glucose. Std1’s predicted strength of repression is greater for transporters with a lower affinity for glucose.

Although the HXTs respond to glucose, we cannot straightforwardly measure its concentration in all these experiments and so instead estimate the concentration from each culture’s optical density (OD). Assuming a constant yield – a fixed amount of glucose must be imported for a cell to replicate, we estimate the glucose concentration as a linear decreasing function of the OD. This function has the culture’s initial glucose concentration at the minimal OD when the experiment begins and is zero when the experiment finishes because the OD has then plateaued (SI).

Building on previous work [4, 6, 32], we draw three conclusions about how the sensors Rtg2 and Snf3 function:

i. The low affinity sensor Rgt2 predominately inactivates the repressor Std1. In glucose (above 0.2%) and when repressed, the low affinity transporters Hxt1 and Hxt3 are repressed almost entirely by Std1 because deleting STD1 but not the other repressor MTH1 substantially increases their levels (Fig. 4A & 4C). This repression by Std1 is relieved in 2% glucose only if Rgt2 is present (Fig. 4A). We observe that Hxt3 is at low levels in 0.2% glucose only if Snf3 is present (Fig. 4C), implying that Snf3 weakly inactivates Std1.
ii. Snf3 predominately inactivates Mth1. Focusing on the medium affinity transporter Hxt2 (Fig. 4B), HXT2 is strongly regulated by Mth1 because deleting MTH1 – and only MTH1 – increases its expression in pyruvate. Hxt2’s levels are reduced more by deleting SNF3 than by deleting RGT2 in 0.2% glucose implying that Snf3 most efficiently inactivates Mth1. There is, however, still some increase in levels of Hxt2 in the *snf3*Δ strain, and so Rgt2 does inactivate Mth1 if weakly. In agreement, HXT6 and HXT7 are also repressed predominately by Mth1 in low glucose and yet partly express if SNF3 is deleted (Fig. 4F & SI).
iii. Mth1 is inactivated at lower concentrations of glucose than Std1. Hxt1 and Hxt3 are absent in pyruvate in a *mthl* Δ and in a *stdl* Δ strain implying that both genes are repressed by Mth1 and Std1 (Fig. 4A & 4C). Yet in 0.2% glucose, Hxt3 levels increase if STD1 is deleted but are unaffected if MTH1 is deleted (Fig. 4C), implying that Mth1 but not Std1 has already been sufficiently inactivated at this concentration of glucose to no longer repress HXT3.

We are also able to infer aspects of each HXT’s transcriptional regulation. As observed in the microfluidic data (Fig. 3C), deleting the repressor STD1 can decrease levels of an Hxt compared to wild-type in sufficiently high glucose and if so deleting RGT2 increases the Hxt’s levels. In batch culture, this phenomenon is most obvious for Hxt4 (Fig. 4D), but occurs too for Hxt2, Hxt6, and Hxt7 (SI) and is absent for Hxt1 and Hxt3 (Fig. 4A & SI). There are two mechanistic explanations (Fig. 3A), which may work together. Std1 inhibits repression by both Mig1 and Mig2 indirectly through promoting SNF1’s activity and by Mig2 directly by repressing the MIG2 gene. Deleting STD1 should therefore reduce levels of an Hxt if its promoter is repressed by Mig1,2. In contrast, deleting RGT2 causes Std1 to hyper-activate increasing its inhibition of Mig1,2’s repression and so raises levels of the Hxt.

Our data therefore support these inferences:

i. HXT1 and HXT3 are repressed by Std1 and by Mth1 in sufficiently low glucose, but only weakly if at all by Mig1 and Mig2.
ii. HXT2, HXT6, and HXT7 are repressed predominantly by Mth1 in low glucose and by Mig1 and Mig2 in high glucose. Repression by Std1 is weak at best.
iii. HXT6 and HXT7 are likely repressed by other regulators in the absence of glucose, such as Msn2 [53] – the master transcription factor for the general stress response, because there is only weak expression in pyruvate when MTH1 is deleted (Fig. 4F & SI) and where we also expect no repression by Mig1,2.
iv. In line with the microfluidic data, HXT4 is repressed by Mth1 and Std1, with Mth1 being dominant in a history of low glucose and Std1 being dominant in a history of high glucose, and by Mig1 and Mig2 in high glucose (Fig. 4D & SI).
v. HXT5 is weakly regulated by the Snf3-Rgt2 system [20] because deleting any one of SNF3, RGT2, MTH1, or STD1 leaves Hxt5’s behaviour unchanged (Fig. 4E & SI). HXT5 is unlikely to be redundantly regulated by Mth1 and Std1 because deleting STD1 at glucose concentrations where Mth1 is expected to be inactivated has no phenotype.

Together the data (Figs. 3 & 4) suggest how cells match transporters of the appropriate affinity to the concentration of glucose. Consider three basic types of transporters with either a high, medium, or low affinity for glucose and whose genes are regulated by two types of repressors. The first type – Mth1 and Std1 – repress when extracellular glucose falls; the second type – Mig1 and Mig2 – repress when extracellular glucose rises. Both types respond together in a push-pull manner in proportion to the concentration of glucose. It is known that Mth1, at least, has a graded inactivation [38], and the level of nuclear Mig1 increases proportionally with the glucose concentration when cells are switched into higher glucose [51] and decreases proportionally when cells are switched into lower glucose [31].

For the high affinity transporters, a rudimentary regulation that allows their expression only in low glucose is to be weakly repressed by Mth1 and strongly repressed by Mig1,2. In the absence of glucose, Mth1 prevents expression. In low glucose, transcription activates because of the reduced level of Mth1, its weak binding to the promoter, and the at best low levels of nuclear Mig1,2. In higher glucose, Mig1,2 increase and shut down transcription.

To enable expression only for intermediate concentrations of glucose, the promoters of the medium affinity transporters should, in contrast, have the binding strengths of Mth1 and Mig1,2 reversed. The promoters should be strongly repressed by Mth1 and weakly repressed by Mig1,2. Mth1 prevents expression both in the absence of and in low glucose because it binds the promoters sufficiently strongly even when its levels decrease in low glucose. In intermediate concentrations of glucose, there is expression because levels of Mth1 become too low to bind and Mig1,2, although present in the nucleus, are not at high enough concentrations to repress given their weak binding. At still higher concentrations, Mig1,2 do prevent transcription.

Expressing the low affinity transporters only in high glucose is simpler. Their promoters should be repressed by both Mth1 and Std1, but not by Mig1,2. Mth1 and Std1 prevent transcription together in the absence of and in sufficiently low glucose. Std1 prevents transcription in intermediate concentrations where Mth1 has been mostly degraded, and transcription activates only in glucose high enough to inactivate Std1.

### Mathematical modelling supports push-pull repression

We used mathematical modelling to determine if this proposed regulation, which we know to be consistent with the data in batch culture (Fig. 4A-F), is also consistent with our microfluidic data (Fig. 2B). To do so, we combined the model of regulation parameterised using the experiments with Hxt4 (Fig. 3C) – this model predicts the levels of the repressors as a function of glucose (Fig. 3D) – with models of transcriptional regulation specific to each HXT. The high and medium affinity transporters – Hxt6, Hxt7, and Hxt2 – are assumed to be regulated only by Mth1 and Mig1,2, and the low affinity transporters – Hxt1 and Hxt3 – only by Mth1 and Std1. We inferred parameters specific to an Hxt – dissociation constants of repressors binding to the promoter, Hill numbers, and the rates of synthesis and degradation – and the rest were fixed at their values found for Hxt4.

These models accurately describe the behaviour of the Hxts in rising and falling glucose (SI), and the resulting best-fit parameters recover the expected binding preferences of the repressors (Fig. 4G-I).

Compared to the promoters for the medium affinity transporters, those for the high affinity transporters are more weakly bound by Mth1 (Fig. 4G) and more strongly repressed by Mig1,2 (Fig. 4H), and Std1, the dominant repressor in high glucose, binds tighter to the promoters of transporters with lower affinities for glucose (Fig. 4I). We calculated the maximal strength of repression using the inferred dissociation constants and Hill numbers and the concentrations of transcription factors we expect from our earlier model (Fig. 3D; SI).

As a stronger test of this proposed regulation, we considered every possible structure of the promoter, assuming that at most all of the repressors and at least one bind, leading to 15 models in total (2^4^ - 1), and asked for each HXT which variant of the promoter is best supported by our data. We used again the model of regulation derived for Hxt4, the data of Fig. 2B, and approximate Bayesian computation to compare statistically the validity of each promoter variant for each HXT. These results confirm our understanding (SI): the data support HXT1 being regulated by Mth1 and Std1 only; and HXT2, HXT6, and HXT7 being regulated by Mth1 and Mig1,2 but not Std1. For HXT3, however, although the data are consistent with regulation by Std1 and Mth1, additional repression by Mig1,2 is also supported. Such regulation has indeed been reported [54].

## Discussion

In this study, we combine systems biology and quantitative time-lapse microscopy to determine both how seven glucose transporters in yeast are expressed in changing concentrations of glucose and the mechanism behind this expression, revealing history-dependent behaviour. Using mathematical modelling and approximate Bayesian computation, we integrated this data with our prior knowledge of the glucose-sensing network to uncover a push-pull system comprising two types of repressors. We find that the dynamics of the transporters is more complex than that expected if cells try to maximise their import given the current availability of glucose and suggests potential strategies that yeast has evolved to be competitive in changing environments.

We hypothesised that multiple transporters with different affinities had been selected because transporters that use facilitated diffusion have a rate-affinity tradeoff. This tradeoff causes each Hxt to import faster than any other for a range of glucose concentrations determined by its affinity for glucose (Fig. 1E). Transporters with high, medium, and low affinity are therefore mostly expressed in low, medium, and high concentrations of glucose, but not entirely (Fig. 2A), hinting that the regulation is also tuned to prepare for likely future events. For each affinity – broadly interpreted, two transporters exist (Fig. 1A) suggesting an advantage to higher dosage: more transporters can be expressed from two genes than from one. Consistently, selection in limiting glucose leads to extra copies of genes for the low affinity transporters, and the mutated strain outcompetes the wild-type in sufficiently low glucose [55].

To express the optimal transporter for the concentration of glucose available, cells could dispense with regulation and simultaneously express all the transporters, but this strategy poorly uses intracellular resources. Repressing genes for suboptimal transporters releases resources to other processes [56, 57, 58], and employing endocytosis to degrade any expressed transporters recycles their amino acids back to the common pool [59].

Furthermore, regulating levels of the high affinity transporters can allow cells to prepare better for starvation, at least for nutrient-sensing systems that control a low and a high affinity transporter [3]. As a nutrient’s availability falls, cells may use the drop in flux through the low affinity transporter as a warning to trigger expression of the high affinity transporter. Cells therefore maintain intracellular nutrients long enough to be able to launch a protective programme before extracellular nutrients are depleted [3]. Although the response to falling glucose is likely to be more complex because cells directly sense extracellular glucose, our results provide a mechanism for how such transcriptional regulation occurs, and cells certainly prepare as glucose decreases: for example, they synthesise and store glycogen [60].

The regulation (Fig. 3A) is approximately bipartite, and this division of labour allows the network to match transporters of approximately three different affinities to three levels of extracellular glucose. The higher affinity sensor Snf3 predominately inactivates the repressor Mth1, which together with Mig1 and Mig2 controls expression of the high and medium affinity transporters; the lower affinity sensor Rgt2 predominately inactivates the repressor Std1, which controls expression of the low affinity transporters. History dependence is generated by the slow response times of Mth1 and Std1 relative to those of Mig1 and Mig2, which create windows of weak repression in falling glucose allowing levels of multiple transporters to spike (Fig. 2B & 3D). Mth1’s slower response is helped by Mig1 and Mig2 repressing its gene. Std1 plays multiple roles. Inactivating Std1 not only directly promotes expression of the low affinity transporters but also indirectly represses expression of the medium and high affinity transporters. Repression by Mig1 and Mig2 increases when Std1 inactivates, because active Std1 inhibits Mig1 and Mig2 by promoting SNF1’s activity and represses MIG2.

The four repressors work as a push-pull system. Mth1 and Std1 repress when extracellular glucose falls; Mig1 and Mig2 repress when extracellular glucose rises. In rising glucose, glucose ‘pulls’ repression by Mth1 and Std1 and ‘pushes’ repression by Mig1 and Mig2; in falling glucose, glucose ‘pulls’ repression by Mig1 and Mig2 and ‘pushes’ repression by Mth1 and Std1 (Fig. 5A). By weakly binding Mth1 and strongly binding Mig1 and Mig2, the promoters for the high affinity transporters express only in low glucose. In contrast, by strongly binding Mth1 and weakly binding Mig1 and Mig2, the promoters for the medium affinity transporters express only in intermediate concentrations of glucose. By binding Std1, the promoters for the low affinity transporters express only in high glucose (Fig. 5A). Although also repressed by Mth1, their expression is principally controlled by Std1 because Mth1 inactivates at lower concentrations of glucose.

**Figure 5.**
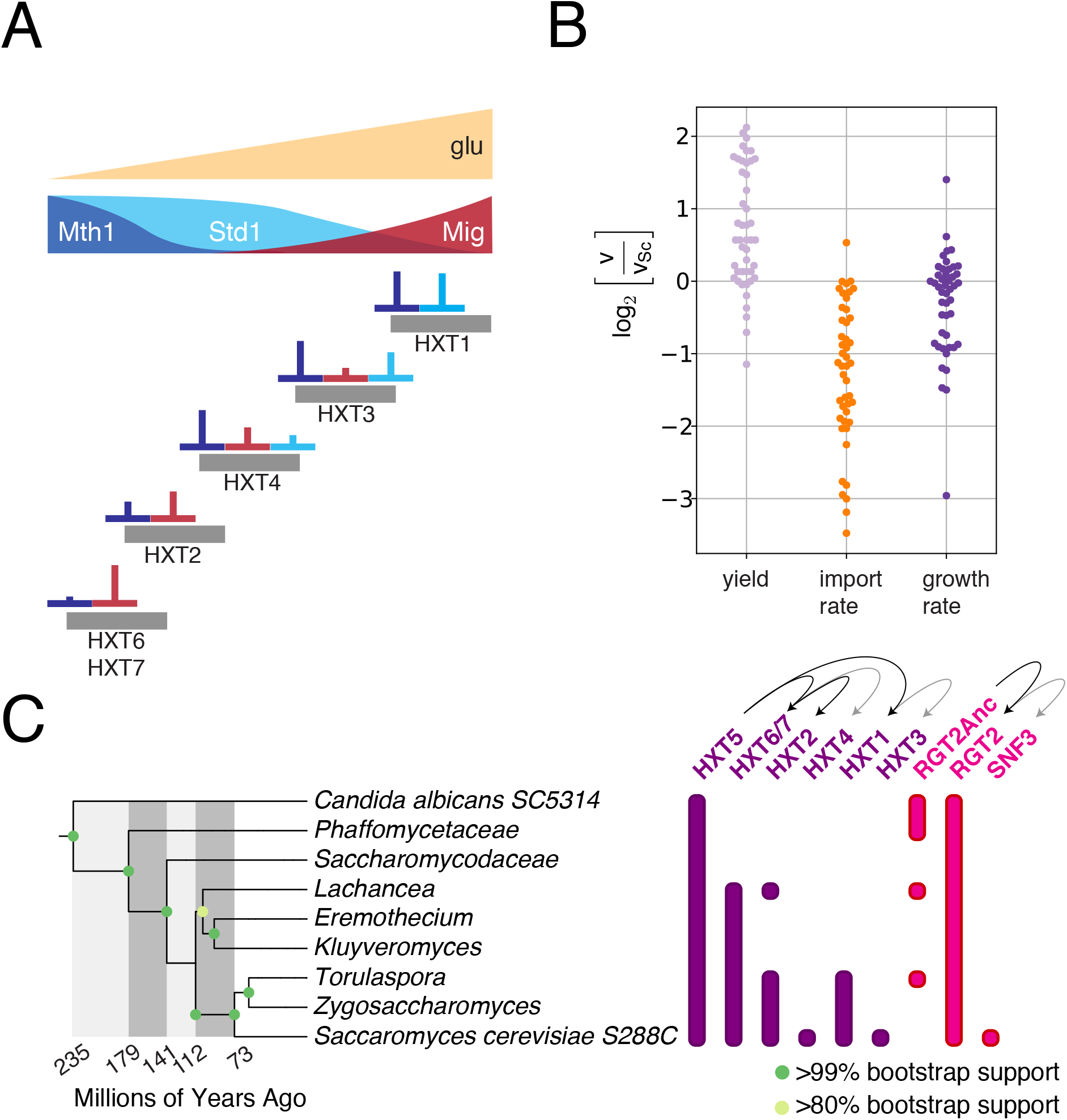
The Hxts are predominately regulated to have high levels in the concentrations of glucose where they import fastest allowing budding yeast to excel at rapidly importing glucose. **A** The push-pull system of repressors allows HXTs to be expressed in the concentrations of glucose that match their affinities for glucose. HXT6 and HXT7 express in low glucose because they bind Mth1 weakly and Mig1,2 strongly; HXT2 and HXT4 express in intermediate glucose because they bind Mth1 strongly and Mig1 and Mig1,2 weakly; HXT1 and HXT3 express in high glucose because they are repressed by Std1 in intermediate and by Std1 and Mth1 in low glucose. HXT3 and HXT4 blur this picture: HXT3 is additionally regulated by Mig1 and HXT4 by Std1. **B** Comparing yield, the rate of glucose import, and growth rate across 46 species of yeast in 2% glucose shows that budding yeast excels at importing glucose (data from [61]). We plot the log_2_ of the ratio of the growth characteristic, denoted *v*, to its value for budding yeast, denoted *v*_Sc_. Negative results imply a lower value than budding yeast. Yield is the amount of biomass produced relative to the amount of glucose consumed. The only species to import at a greater rate that budding yeast is another *Saccharomyces* species – *S. mikatae*, which has 19 HXTs compared to *S. cerevisiae*’s 18 [12]. **C** A phylogenetic analysis suggests that HXT4, HXT3, and SNF3 are relatively recent additions to budding yeast’s genome. The arrows show the likely origins of duplications. RGT2Anc is the ancestor of today’s RGT2.

Although the regulation broadly follows this proposed structure, the categories are blurred. HXT3 is also repressed by Mig1 [54] (SI) and HXT4 by Std1 (Fig. 3B & 4D). There is additional regulation too to integrate glucose transport with other cellular responses. For example, the transcription of HXT1 is promoted by the kinase Hog1 in hyperosmotic stress [62] and the transcription of HXT4 by the transcription factors regulating glycolytic genes [63]. HXT2 too is special. Its mRNA – but not those from the other HXTs – is enriched in the buds that form when glucose is added to starving cells [64].

A possible advantage of the system responding to metabolised intracellular glucose through Mig1 and Mig2, as well as to extracellular glucose through Mth1 and Std1, is that transport changes ‘gears’ from high to medium affinity and from medium to low affinity and vice versa only if the sensing of extracellular glucose is matched with the level of metabolised glucose (Fig. 3A). This precaution may limit costs from switching metabolism needlessly between fermentation and respiration.

An exception to the regulation is Hxt5, which appears to be at best weakly regulated by the Snf3-Rgt2 network (Fig. 4E) and is expressed only in sufficiently low glucose despite being a transporter of medium affinity (Fig. 1A & 2B). We suspect that Hxt5 is necessary to prime the response because the other HXTs are regulated by metabolised glucose. If starving cells are exposed to high or medium levels of glucose, then Snf3 and Rgt2 will cause both the high and medium affinity transporters to be expressed. Only when these transporters have enabled sufficient metabolism of glucose will Mig1,2 activate enough to repress the genes for the inefficient high affinity transporters. Expressing and then repressing these transporters is both metabolically wasteful and lengthens the time for the cells to generate the glycolytic flux appropriate to the sugar available. With a priming transporter that enables immediate catabolism of glucose, however, Mig1,2 may be activated quicker and so prevent the levels of the high affinity transporters ever becoming substantial. This priming transporter should be of medium affinity, like Hxt5, to prevent expression of the high affinity transporters in medium and high glucose, and it should be expressed when the other HXTs are not with its own distinct regulation. Consistent with a priming function, mutants missing HXT5 grow more slowly than the wild type if 2% glucose is given to starving cells [30, 65].

Although we have focused on transcriptional regulation, it is not the cell’s only response. Levels of the Hxts are also tuned by active degradation, which is controlled by the kinases PKA and SNF1. In high glucose, PKA inhibits the endocytosis of the low affinity transporters [66, 67]. In falling glucose, the medium and high affinity transporters are degraded via the arrestin-related trafficking adaptor Csr2, whose expression is controlled by SNF1 through Mig1 and Mig2 and whose activity is also inhibited by PKA [52]. In rising glucose, however, a different adaptor Rod1 is required, at least for Hxt6 [68]. Such regulation might target inactive transporters allowing the cell to correct inappropriate matching of transporters to the concentration of glucose available.

We suspect that the HXT network has been selected to make budding yeast competitive through its ability to sequester glucose from other species. The number of HXTs together with the regulation to ensure that import is dominated by the optimal transporter for the current concentration of glucose enables an exceptionally rapid uptake (Fig. 5B). Competition becomes most fierce when a resource becomes scarce, and it is in falling glucose that we observe spikes of expression – likely to capture whatever glucose remains, first for the medium affinity transporters and then for the high affinity ones (Fig. 2B). Even in constant concentrations, cells express suboptimal transporters, although to lower levels than the optimal one (Fig. 2A), and are potentially poised to respond if the concentration falls – a bet-hedging strategy where a lower fitness in the current environment is traded for a fast response and so a higher fitness in a future one.

Cells would not necessarily be selected to metabolise glucose efficiently but to starve competitors [69], and such selfish strategies have been predicted to be evolutionarily stable [70]. Budding yeast do ferment rather than respire glucose in sufficiently high concentrations despite then generating substantially less ATP per glucose molecule. This choice may allow cells to have a glycolytic flux higher than that which can be processed by mitochondria [71] and across yeast species correlates with having high numbers of genes for hexose transporters [12, 72]. To consume the ethanol generated by this fermentation, cells must respire, which involves a substantial re-routing of metabolism [73] and causes growth to lag [74, 75] and even to arrest [75, 76]. An alternative explanation for the spikes of expression of Hxts we observe in falling glucose (Fig. 2B) is therefore to maintain fermentative growth for as low concentrations of glucose as possible so as to delay this costly metabolic transition. Ethanol is indeed only consumed in batch cultures when glucose is completely exhausted [61].

Both the sensors SNF3 and RGT2 and the repressors MTH1 and STD1 were created when budding yeast’s whole genome was duplicated [77]. The duplication allowed the two sensors to specialise. SNF3 evolved to sense glucose with high affinity while regulating the high and medium affinity transporters via Mth1. RGT2 specialised to sensing glucose with low affinity and so allowed the evolution of fast low affinity transporters – potentially the last to evolve in budding yeast – with their own distinct regulation dominated by Std1 (Fig. 5C).

Methodologically, our work illustrates a general integrative approach to unravelling cellular regulation. First, we used quantitative time-series experiments with both informative timevarying inputs and a range of genetic perturbations to gather sufficiently rich data. Second, we integrated this data with prior knowledge of the network’s structure using mathematical modelling and Bayesian methods of model comparison. Third, we used a modular approach: performing initially a broad analysis focused on one output of the system to characterise the regulatory module and then more focused experiments to determining quantitatively how the other outputs are regulated given that regulatory module.

The task of matching expression of a gene to a particular pattern of input is common [1], and the solution exhibited by the HXT network may be a convergent one. The network enables a transporter to be expressed in a restricted range of glucose concentrations by having two types of repressors whose level of repression changes with the concentration of glucose in opposite ways – one type increasing as glucose rises and the other decreasing in a push-pull system. This behaviour mirrors the regulation acting in a different setting – the early development of the *Drosophila* embryo [2]. There the task is, broadly, to match expression of developmental genes to a restricted spatial range [78], and a key component of the solution is having the activity of transcription factors change with space in opposite ways – the concentration of some transcription factors increasing with distance from the embryo’s anterior pole and that of others decreasing. We expect that this regulatory strategy is widespread.

## Acknowledgements

We thank Dr Ivan Clark for his technical assistance and advice and gratefully acknowledge funding by the Wellcome Trust (LFMG) and the BBSRC (LFMG, VS, and PSS).

## References

[1] Jacob, F. & Monod, J. Genetic regulatory mechanisms in the synthesis of proteins. J Mol Biol 3, 318–356 (1961).

[2] St Johnston, D. & Nüsslein-Volhard, C. The origin of pattern and polarity in the Drosophila embryo. Cell 68, 201–219 (1992).

[3] Levy, S., Kafri, M., Carmi, M. & Barkai, N. The competitive advantage of a dual-transporter system. Science 334, 1408–1412 (2011).

[4] Bisson, L. F., Fan, Q. & Walker, G. A. Sugar and glycerol transport in Saccharomyces cerevisiae. In Yeast Membrane Transport, 125–168 (Springer, 2016).

[5] Wieczorke, R. et al. Concurrent knock-out of at least 20 transporter genes is required to block uptake of hexoses in Saccharomyces cerevisiae. FEBS Letters 464, 123–128 (1999).

[6] Ozcan, S. & Johnston, M. Three different regulatory mechanisms enable yeast hexose transporter (HXT) genes to be induced by different levels of glucose. Mol Cell Biol 15, 1564–1572 (1995).

[7] Klockow, C., Stahl, F., Scheper, T. & Hitzmann, B. In vivo regulation of glucose transporter genes at glucose concentrations between 0 and 500 mg/L in a wild type of Saccharomyces cerevisiae. J Biotechn 135, 161–167 (2008).

[8] Youk, H. & van Oudenaarden, A. Growth landscape formed by perception and import of glucose in yeast. Nature 462, 875–879 (2009).

[9] Zaman, S., Lippman, S. I., Schneper, L., Slonim, N. & Broach, J. R. Glucose regulates transcription in yeast through a network of signaling pathways. Mol Syst Biol 5, 245 (2009).

[10] Marinkovic, Z. S. et al. A microfluidic device for inferring metabolic landscapes in yeast monolayer colonies. Elife 8, e47951 (2019).

[11] Maier, A., Volker, B., Boles, E. & Fuhrmann, G. F. Characterisation of glucose transport in Saccharomyces cerevisiae with plasma membrane vesicles (countertransport) and intact cells (initial uptake) with single Hxt1, Hxt2, Hxt3, Hxt4, Hxt6, Hxt7 or Gal2 transporters. FEMS Yeast Res 2, 539–550 (2002).

[12] Lin, Z. & Li, W.-H. Expansion of hexose transporter genes was associated with the evolution of aerobic fermentation in yeasts. Mol Biol Evol 28, 131–142 (2011).

[13] Özcan, S. & Johnston, M. Function and regulation of yeast hexose transporters. Microbiol Mol Biol Rev 63, 554–569 (1999).

[14] Broach, J. R. Nutritional control of growth and development in yeast. Genetics 192, 73–105 (2012).

[15] Gudelj, I., Beardmore, R. E., Arkin, S. & MacLean, R. C. Constraints on microbial metabolism drive evolutionary diversification in homogeneous environments. J Evol Biol 20, 1882–1889 (2007).

[16] Bisson, L. F., Coons, D. M., Kruckeberg, A. L. & Lewis, D. A. Yeast sugar transporters. Crit Rev Biochem Mol Biol 28, 259–308 (1993).

[17] Reifenberger, E., Boles, E. & Ciriacy, M. Kinetic characterization of individual hexose transporters of Saccharomyces cerevisiae and their relation to the triggering mechanisms of glucose repression. Eur J Biochem 245, 324–333 (1997).

[18] Wolfe, K. H. & Shields, D. C. Molecular evidence for an ancient duplication of the entire yeast genome. Nature 387, 708–713 (1997).

[19] Verwaal, R. et al. HXT5 expression is determined by growth rates in Saccharomyces cerevisiae. Yeast 19, 1029–1038 (2002).

[20] Verwaal, R. et al. HXT5 expression is under control of STRE and HAP elements in the HXT5 promoter. Yeast 21, 747–757 (2004).

[21] van Suylekom, D. et al. Degradation of the hexose transporter Hxt5p in Saccharomyces cerevisiae. Biol Cell 99, 13–23 (2007).

[22] Fersht, A. Structure and mechanism in protein science (WH Freeman, New York, 1999).

[23] Teusink, B., Diderich, J. A., Westerhoff, H. V., Van Dam, K. & Walsh, M. C. Intracellular glucose concentration in derepressed yeast cells consuming glucose is high enough to reduce the glucose transport rate by 50%. J Bacteriol 180, 556–562 (1998).

[24] Bosdriesz, E. et al. Low affinity uniporter carrier proteins can increase net substrate uptake rate by reducing efflux. Sci Rep 8, 5576 (2018).

[25] Crane, M. M., Clark, I. B., Bakker, E., Smith, S. & Swain, P. S. A microfluidic system for studying ageing and dynamic single-cell responses in budding yeast. PLoS One 9, e100042 (2014).

[26] Granados, A. A. et al. Distributing tasks via multiple input pathways increases cellular survival in stress. eLife 6, e21415 (2017).

[27] Bakker, E., Swain, P. S. & Crane, M. M. Morphologically constrained and data informed cell segmentation of budding yeast. Bioinformatics 34, 88–96 (2017).

[28] Ronen, M. & Botstein, D. Transcriptional response of steady-state yeast cultures to transient perturbations in carbon source. Proc Nat Acad Sci USA 103, 389–394 (2006).

[29] Levy, S. & Barkai, N. Coordination of gene expression with growth rate: a feedback or a feed-forward strategy? FEBS Lett 583, 3974–3978 (2009).

[30] Diderich, J. A., Merijn Schuurmans, J., Van Gaalen, M. C., Kruckeberg, A. L. & Van Dam, K. Functional analysis of the hexose transporter homologue HXT5 in Saccharomyces cerevisiae. Yeast 18, 1515–1524 (2001).

[31] Granados, A. A. et al. Distributed and dynamic intracellular organization of extracellular information. Proc Nat Acad Sci USA 115, 6088–6093 (2018).

[32] Ozcan, S., Dover, J., Rosenwald, A. G., Wölfl, S. & Johnston, M. Two glucose transporters in Saccharomyces cerevisiae are glucose sensors that generate a signal for induction of gene expression. Proc Nat Acad Sci USA 93, 12428–12432 (1996).

[33] Lakshmanan, J., Mosley, A. L. & Özcan, S. Repression of transcription by Rgt1 in the absence of glucose requires Std1 and Mth1. Curr Genet 44, 19–25 (2003).

[34] Polish, J. A., Kim, J.-H. & Johnston, M. How the Rgt1 transcription factor of Saccharomyces cerevisiae is regulated by glucose. Genetics 169, 583–594 (2005).

[35] Snowdon, C. & Johnston, M. A novel role for yeast casein kinases in glucose sensing and signaling. Mol Biol Cell 27, 3369–3375 (2016).

[36] Flick, K. M. et al. Grr1-dependent inactivation of Mth1 mediates glucose-induced dissociation of Rgt1 from HXT gene promoters. Mol Biol Cell 14, 3230–3241 (2003).

[37] Kaniak, A., Xue, Z., Macool, D., Kim, J.-H. & Johnston, M. Regulatory network connecting two glucose signal transduction pathways in Saccharomyces cerevisiae. Eukaryot Cell 3, 221–231 (2004).

[38] Sabina, J. & Johnston, M. Asymmetric signal transduction through paralogs that comprise a genetic switch for sugar sensing in Saccharomyces cerevisiae. J Biol Chem 284, 29635–29643 (2009).

[39] Simpson-Lavy, K., Xu, T., Johnston, M. & Kupiec, M. The Std1 activator of the Snf1/AMPK kinase controls glucose response in yeast by a regulated protein aggregation. Mol Cell 68, 1120–1133 (2017).

[40] Mayer, F. V. et al. ADP regulates SNF1, the Saccharomyces cerevisiae homolog of AMP-activated protein kinase. Cell Metab 14, 707–714 (2011).

[41] Treitel, M. A., Kuchin, S. & Carlson, M. Snf1 protein kinase regulates phosphorylation of the Mig1 repressor in Saccharomyces cerevisiae. Mol Cell Biol 18, 6273–6280 (1998).

[42] Lim, M. K. et al. Galactose induction of the GAL1 gene requires conditional degradation of the Mig2 repressor. Biochem J 435, 641–649 (2011).

[43] Hubbard, E., Jiang, R. & Carlson, M. Dosage-dependent modulation of glucose repression by MSN3 (STD1) in Saccharomyces cerevisiae. Mol Cell Biol 14, 1972–1978 (1994).

[44] Tomás-Cobos, L. & Sanz, P. Active Snf1 protein kinase inhibits expression of the Saccharomyces cerevisiae HXT1 glucose transporter gene. Biochem J 368, 657 (2002).

[45] Kuchin, S., Vyas, V. K., Kanter, E., Hong, S.-P. & Carlson, M. Std1p (Msn3p) positively regulates the Snf1 kinase in Saccharomyces cerevisiae. Genetics 163, 507–514 (2003).

[46] Jouandot, D., Roy, A. & Kim, J.-H. Functional dissection of the glucose signaling pathways that regulate the yeast glucose transporter gene (HXT) repressor Rgt1. J Cell Biochem 112, 3268–3275 (2011).

[47] Westholm, J. O. et al. Combinatorial control of gene expression by the three yeast repressors mig1, mig2 and mig3. BMC Genomics 9, 601 (2008).

[48] Schmidt, M. C. et al. Std1 and Mth1 proteins interact with the glucose sensors to control glucose-regulated gene expression in Saccharomyces cerevisiae. Mol Cell Biol 19, 4561–4571 (1999).

[49] Dietzel, K. L., Ramakrishnan, V., Murphy, E. E. & Bisson, L. F. MTH1 and RGT1 demonstrate combined haploinsufficiency in regulation of the hexose transporter genes in Saccharomyces cerevisiae. BMC Genet 13, 1–10 (2012).

[50] Bisson, L., Neigeborn, L., Carlson, M. & Fraenkel, D. The SNF3 gene is required for high-affinity glucose transport in Saccharomyces cerevisiae. J Bacteriol 169, 1656–1662 (1987).

[51] Bendrioua, L. et al. Yeast AMP-activated protein kinase monitors glucose concentration changes and absolute glucose levels. Journal of Biological Chemistry 289, 12863–12875 (2014).

[52] Hovsepian, J. et al. Multilevel regulation of an *α*-arrestin by glucose depletion controls hexose transporter endocytosis. J Cell Biol 216, 1811–1831 (2017).

[53] Pfanzagl, V. et al. A constitutive active allele of the transcription factor Msn2 mimicking low PKA activity dictates metabolic remodeling in yeast. Mol Biol Cell 29, 2848–2862 (2018).

[54] Kuttykrishnan, S., Sabina, J., Langton, L. L., Johnston, M. & Brent, M. R. A quantitative model of glucose signaling in yeast reveals an incoherent feed forward loop leading to a specific, transient pulse of transcription. Proc Nat Acad Sci USA 107, 16743–16748 (2010).

[55] Brown, C. J., Todd, K. M. & Rosenzweig, R. F. Multiple duplications of yeast hexose transport genes in response to selection in a glucose-limited environment. Mol Biol Evol 15, 931–942 (1998).

[56] Stoebel, D. M., Dean, A. M. & Dykhuizen, D. E. The cost of expression of Escherichia coli lac operon proteins is in the process, not in the products. Genetics 178, 1653–1660 (2008).

[57] Lang, G. I., Murray, A. W. & Botstein, D. The cost of gene expression underlies a fitness trade-off in yeast. Proc Nat Acad Sci USA 106, 5755–5760 (2009).

[58] Kafri, M., Metzl-Raz, E., Jona, G. & Barkai, N. The cost of protein production. Cell Rep 14, 22–31 (2016).

[59] Goode, B. L., Eskin, J. A. & Wendland, B. Actin and endocytosis in budding yeast. Genetics 199, 315–358 (2015).

[60] Lillie, S. H. & Pringle, J. R. Reserve carbohydrate metabolism in Saccharomyces cerevisiae: responses to nutrient limitation. J Bacteriol 143, 1384–1394 (1980).

[61] Hagman, A., Säll, T., Compagno, C. & Piskur, J. Yeast “make-accumulate-consume” life strategy evolved as a multi-step process that predates the whole genome duplication. PLoS One 8, e68734 (2013).

[62] Tomás-Cobos, L., Casadomé, L., Mas, G., Sanz, P. & Posas, F. Expression of the HXT1 low affinity glucose transporter requires the coordinated activities of the HOG and glucose signalling pathways. J Biol Chem 279, 22010–22019 (2004).

[63] Türkel, S. & Bisson, L. F. Transcription of the HXT4 gene is regulated by Gcr1p and Gcr2p in the yeast S. cerevisiae. Yeast 15, 1045–1057 (1999).

[64] Stahl, T. et al. Asymmetric distribution of glucose transporter mRNA provides a growth advantage in yeast. EMBO J 38, e100373 (2019).

[65] Bermejo, C., Haerizadeh, F., Takanaga, H., Chermak, D. & Frommer, W. B. Dynamic analysis of cytosolic glucose and ATP levels in yeast using optical sensors. Biochem J 432, 399–406 (2010).

[66] Snowdon, C. & Van der Merwe, G. Regulation of Hxt3 and Hxt7 turnover converges on the Vid30 complex and requires inactivation of the Ras/cAMP/PKA pathway in Saccharomyces cerevisiae. PLoS One 7, e50458 (2012).

[67] Roy, A., Kim, Y.-B., Cho, K. H. & Kim, J.-H. Glucose starvation-induced turnover of the yeast glucose transporter Hxt1. Biochim Biophys Acta 1840, 2878–2885 (2014).

[68] Nikko, E. & Pelham, H. R. Arrestin-mediated endocytosis of yeast plasma membrane transporters. Traffic 10, 1856–1867 (2009).

[69] Hagman, A. & Piŝkur, J. A study on the fundamental mechanism and the evolutionary driving forces behind aerobic fermentation in yeast. PLoS One 10, e0116942 (2015).

[70] Josephides, C. & Swain, P. S. Predicting metabolic adaptation from networks of mutational paths. Nat Commun 8, 685 (2017).

[71] Sonnleitner, B. & Käppeli, O. Growth of Saccharomyces cerevisiae is controlled by its limited respiratory capacity: formulation and verification of a hypothesis. Biotechnol Bioeng 28, 927–937 (1986).

[72] Conant, G. C. & Wolfe, K. H. Increased glycolytic flux as an outcome of whole-genome duplication in yeast. Mol Syst Biol 3, 129 (2007).

[73] Frick, O. & Wittmann, C. Characterization of the metabolic shift between oxidative and fermentative growth in Saccharomyces cerevisiae by comparative 13 C flux analysis. Microb Cell Fact 4, 1–16 (2005).

[74] Cerulus, B. et al. Transition between fermentation and respiration determines history-dependent behavior in fluctuating carbon sources. Elife 7, e39234 (2018).

[75] Bagamery, L. E., Justman, Q. A., Garner, E. C. & Murray, A. W. A putative bet-hedging strategy buffers budding yeast against environmental instability. Curr Biol 30, 4563–4578 (2020).

[76] Jacquel, B., Aspert, T., Laporte, D., Sagot, I. & Charvin, G. pH fluctuations drive waves of stereotypical cellular reorganizations during entry into quiescence. bioRxiv (2020).

[77] Byrne, K. P. & Wolfe, K. H. The Yeast Gene Order Browser: combining curated homology and syntenic context reveals gene fate in polyploid species. Genome Res 15, 1456–1461 (2005).

[78] Dubuis, J. O., Tkačik, G., Wieschaus, E. F., Gregor, T. & Bialek, W. Positional information, in bits. Proc Nat Acad Sci USA 110, 16301–16308 (2013).

